# Beta and theta oscillations differentially support free versus forced control over multiple-target search

**DOI:** 10.1101/422691

**Authors:** Joram van Driel, Eduard Ort, Johannes J. Fahrenfort, Christian N. L. Olivers

**Author notes:** Shared senior authorship (Johannes J. Fahrenfort, Christian N. L. Olivers). **Corresponding author**: Christian N.L. Olivers, Department of Experimental and Applied Psychology, Institute for Brain and Behavior, Vrije Universiteit, Amsterdam. Van der Boechorststraat 7, 1081 BT Amsterdam, The Netherlands.

## Abstract

Many important situations require human observers to simultaneously search for more than one object. Despite a long history of research into visual search, the behavioral and neural mechanisms associated with multiple-target search are poorly understood. Here we test the novel theory that the efficiency of looking for multiple targets critically depends on the mode of cognitive control the environment affords to the observer. We used an innovative combination of EEG and eye tracking while participants searched for two targets, within two different contexts: Either both targets were present in the search display and observers were free to prioritize either one of them, thus enabling proactive control over selection; or only one of the two targets would be present in each search display, which requires reactive control to reconfigure selection when the wrong target is prioritized. During proactive control, both univariate and multivariate signals of beta-band (15–35 Hz) power suppression prior to display onset predicted switches between target selections. This signal originated over midfrontal and sensorimotor regions and has previously been associated with endogenous state changes. In contrast, imposed target selections requiring reactive control elicited prefrontal power enhancements in the delta/theta-band (2–8 Hz), but only after display onset. This signal predicted individual differences in associated oculomotor switch costs, reflecting reactive reconfiguration of target selection. The results provide compelling evidence that multiple target representations are differentially prioritized during visual search, and for the first time reveal distinct neural mechanisms underlying proactive and reactive control over multiple-target search.

**Significance Statement:** Searching for more than one object in complex visual scenes can be detrimental for search performance. While perhaps annoying in daily life, this can have severe consequences in professional settings such as medical and security screening. Previous research has not yet resolved whether multiple-target search involves changing priorities in what people attend to, and how such changes are controlled. We approached these questions by concurrently measuring cortical activity and eye movements using EEG and eye tracking, while observers searched for multiple possible targets. Our findings provide the first unequivocal support for the existence of two modes of control during multiple-target search, which are expressed in qualitatively distinct time-frequency signatures of the EEG both before and after visual selection.

---

Baggage scanning, medical image screening, and sports match refereeing are just a few of the activities in which human observers are required to look for multiple relevant visual signals. Studies of visual search behavior have found that multiple-target search comes with performance costs (1-7). Targets are detected more slowly, and are more often missed when observers try to search for more than a single object simultaneously. These costs emerge in particular when the target changes between consecutive searches, compared to when it repeats. Such switch costs suggest changes in the prioritized state of one target representation over others (8, 9), with potentially just a single target representation being activated at any moment in time (10). However, other studies have reported evidence that two different target objects can be found interchangeably without switch costs, thus supporting theories which state that multiple targets can be prioritized equally and in parallel (11-14).

Here we address the question whether multiple-target search indeed involves switch-related changes in target priority state, and if so, how such state changes are controlled. Recent behavioral evidence from our lab indicates that the environmental context, and the type of cognitive control mechanisms it allows for, determine the occurrence of switch costs in multiple-target search (15, 16). Using a gaze-contingent search task in which observers looked for two different targets, we found that saccade latencies were prolonged when the target changed from one trial to the next, but only so when either one of the targets was available per display. When *both* sought-for targets were available in each display, observers still frequently switched from one target to the other, but now without switch costs. This dependence of switch costs on target availability can be explained by assuming that a) at the start of each trial one target representation is prioritized over the other, and b) this prioritization calls for different cognitive control mechanisms. Specifically, when only one target stimulus is available per display, target switches are necessarily imposed upon the observer. If the wrong target happens to be prioritized, this requires a reactive reconfiguration in order to select the unanticipated target (cf. (8, 17)). By definition, such reactive processes can only start *after* display onset, resulting in time costs.

In contrast, when both targets are available in the display, there is no wrong choice. Observers can freely choose which target to select next, and can thus switch latently, *prior* to each display, with little to no switch cost as a result. The difference between enforced, reactive control and free, proactive control has been proposed before in the context of task switches (18), distractor suppression (19), and spatial cueing (20), but its role in visual search is currently unknown.

Importantly, the cognitive control account claims that target switches are accompanied by changes in cognitive state, as priority settings are being reconfigured from one target representation to the other. In contrast, theories that claim that multiple target representations can be prioritized equally and concurrently do not need to assume such switch-related control processes, because no priority changes are required in the first place. Because eye movements are only the end result of the selection process, the existing oculomotor data cannot distinguish between these accounts, and a more direct measure of cognitive states both before and after target switches is required. We therefore used a hybrid approach of concurrently measuring both eye gaze and the electroencephalogram (EEG) of participants instructed to look simultaneously for two different color-defined targets. This gave use the unique opportunity to relate ongoing electrophysiological brain dynamics to any potential switch-related control processes surrounding saccadic target selection. Specifically, we tested the hypothesis that free target choice during search is supported by endogenously triggered, proactive control that may be akin to internally driven, voluntary action selection (21, 22), and most likely originates in medial and lateral frontal cortical areas (20, 23, 24). Crucially, the signal reflecting this control should already emerge *before* a target switch. Conversely, we hypothesized any reactive control-related signal changes to occur only *after* a target switch, specifically when such switches are imposed. Reactive control has previously been tied to an increase in oscillatory power in the theta frequency range (3–8 Hz) over prefrontal brain areas after an unexpected task-switch (25), a novel stimulus (26), or response conflict (27), but its role in visual target selection is unknown.

## Results

A planned number of 30 participants performed a gaze-contingent memory-guided visual search task (Fig 1A). At the start of every sequence of trials, observers were given a cue as to which two target colors to look for. This was followed by 40 consecutive search displays each containing a heterogeneous set of colors, among which either one or two target colors (depending on condition). Participants were instructed to make an eye movement towards one of the two target colors, while avoiding distractors. A new display would then emerge with the current fixation as the starting point. Crucially, in the *Free selection* condition, both target colors were always present in each search display, thus allowing observers proactive control over which target to prioritize from trial to trial. In the *Imposed selection* condition, each search display only contained one of the two target colors, and which target would appear was randomly determined (with the same distribution of switches as in the Free selection condition; see Methods). Thus here, prioritization for the wrong target would require reactive priority reconfiguration. However, if instead observers prioritize both targets equally, no changes in proactive nor reactive control are necessary from trial to trial.

**Fig. 1.**
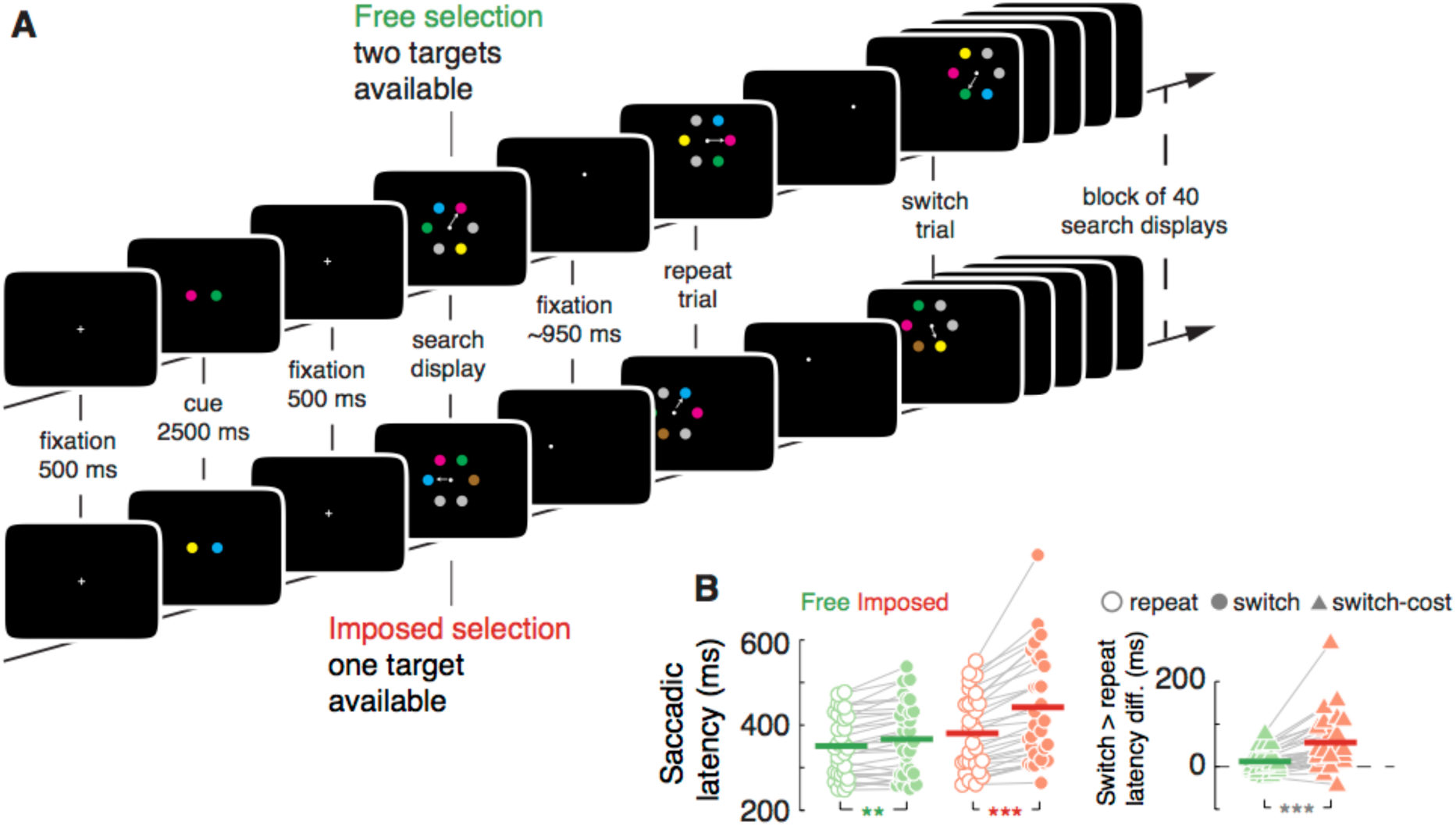
Task design and behavioral results. A) A block began with a fixation cross, and a cue indicating the two target colors for the subsequent sequence of search displays. Depending on the condition each search display contained either one target color (Imposed selection condition; hypothesized to require reactive control on a significant portion of trials) or both target colors (Free selection condition; allowing for efficient proactive control throughout the block). Participants were required to make an eye-movement to, and fixate (one of) the (two) target(s), which then triggered the next display. B) Left: Saccadic latency in ms as a function of condition (Green: Free selection condition; Red: Imposed selection condition) and trial type (open dots: repeat trials; filled dots: switch trials). Each dot shows the trial-average data of a single observer. Horizontal lines show the group-average. Grey lines connecting the dots visualize the within-subject difference between repeat and switch trials. Colored asterisks show the within-condition comparison of repeat versus switch trials, thus illustrating switch costs in both conditions. Right: Switch-costs in ms for Free selection (green triangles) and Imposed selection (red triangles) conditions. Grey lines and asterisk show the interaction effect of a stronger switch costs in the Imposed selection than in the Free selection condition. ^∗∗^ p < 0.01; ^∗∗∗^ p < 0.001.

Our results clearly show differentially controlled priority states. First, switch costs were considerably larger in the Imposed than in the Free selection condition (*F*_1,29_ = 14.66, *p* < 0.001; Fig. 1B). When a change in targets was task-imposed, observers were slower than when targets repeated from one trial to the next (*t*_29_ = 5.20, *p* < 0.001). In fact, there was also a reliable, though much smaller, switch cost when target selection was free (*t*_29_ = 3.36, *p* = 0.002). These magnitude differences in switch costs are a direct replication of earlier findings (15, 16), and are consistent with, though not conclusive for, a difference in the latency and type of control.

Second, the EEG data reveal clear differential state changes associated with freely initiated versus task-imposed switches. In the Free selection condition, any neural signature reflecting a preparatory mechanism should be apparent before display onset, in the time between the offset of the previous and the onset of the next search display. We moreover hypothesized a potential proactive control mechanism to show a topographical distribution over midfrontal scalp regions, consistent with neuroimaging studies on voluntary and self-initiated behavior (23, 24). Local oscillatory dynamics have been linked to memory content, motor intentions, and different modes of control (28, 29). Therefore, we decomposed the EEG data into its time-frequency representation, and compared switch-related activity in three frontocentral electrodes (FC1, FCz and FC2) for Free versus Imposed selection. The Free selection condition showed a robust reduction of power in the beta-band (15-35 Hz) for switch relative to repeat trials (cluster-corrected *p* < 0.001; Fig. 2A), starting ~700 ms prior to the upcoming search display. The effect comprised one sustained time-frequency cluster, reducing in bandwidth around the moment of the saccade, after which the same broadband beta suppression effect re-emerged postsaccade. Importantly, this effect was not present in the Imposed selection condition (*p* > 0.90). This difference was also apparent from the Free versus Imposed selection contrast, which showed significant switch-related beta-suppression in a time window ~500 ms prior to the anticipated onset of the next search display (*p* = 0.002), followed by post-stimulus (and peri-saccade) suppression in the upper-alpha/lower-beta range (10–18 Hz; ~200–900 ms; *p* = 0.002). This shows that in a context of free choice, proactively deciding to switch to a different target is supported by relatively suppressed midfrontal beta power.

**Fig. 2.**
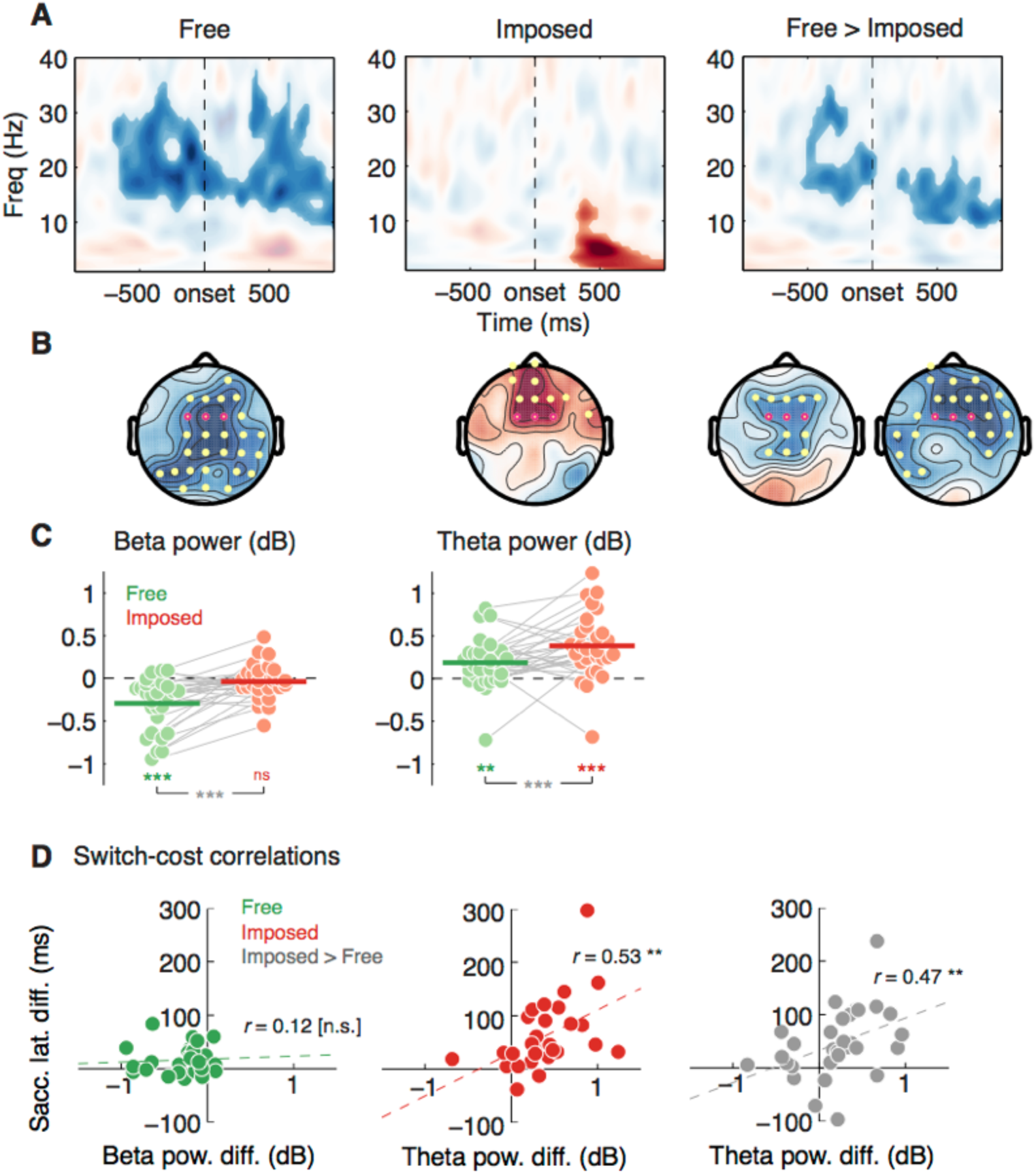
Local time-frequency power results. A) Time-frequency maps showing t-values of statistical tests of switch>repeat on decibel-normalized power, over three midfrontal channels (FC1, FCz, FC2; highlighted in purple in the topographical maps in B, for the Free and Imposed selection condition, and their contrast. Saturated colors show clusters of contiguously significant (p < 0.05) time-frequency points after correcting for cluster sizes under the null-hypothesis of no effect (cluster-threshold: p < 0.05). B) Topographical map of t-values, averaged over the significant time-frequency clusters shown in A. Purple disks show the three midfrontal channels of the time-frequency map, yellow disks show the extension of the effects shown after testing over all channels and using a cluster-size threshold of p < 0.01. C) Single-subject power (dB) for switch>repeat, averaged within the beta-band (left) and theta-band (right) time-frequency clusters shown in A. Green dots: Free selection condition; red dots: Imposed selection condition. Horizontal lines show the group-average. Grey lines connecting the dots visualize the within-subject difference between Free and Imposed selection; ^∗∗^ p < 0.01; ^∗∗∗^ p < 0.001. D) Scatter plots showing across-subject the spectral power of switch versus repeat trials, against behavioral switch costs (saccadic-latency difference of switch versus repeat trials), per condition (left and center), and for the condition difference of Free versus Imposed (right). ^∗∗^ significant robust percentage-bend correlation (30), p < 0.01.

To further test the putative involvement of frontal control regions, we evaluated the topographical specificity of these effects. We averaged the activity within the beta-band time-frequency windows, and tested for clusters of channels that would show differences between switch and repeat trials and between conditions. This revealed that the beta-band modulation in the Free selection condition covered prefrontal as well as posterior parietal scalp regions, with a right-hemisphere dominance (*p* < 0.001; Fig. 2B), while there was again no effect in the Imposed selection condition. The pre-stimulus difference between conditions was localized to a more confined midfrontal-premotor region (*p* < 0.001). Thus we link the endogenous switching between target representations to modulations of frontoparietal beta oscillations.

In contrast, switches in the Imposed selection condition elicited robust low-frequency (2–8 Hz) delta-to-theta band power enhancements starting ~250 ms post-display onset, over the three midfrontal channels (Fig. 2A; *p* = 0.002), which topographically extended towards anterior and lateral prefrontal scalp regions (*p* < 0.001; Fig. 2B). Similar medial and lateral prefrontal theta-band modulations have been linked to a myriad of conflict- and error-related performance monitoring mechanisms (for reviews see (27, 31)). When we tested within the theta-band cluster only, both conditions showed switch-related theta-power enhancement (Free: *t*_29_ = 3.49, *p* = 0.002; Imposed: *t*_29_ = 5.54, *p* < 0.001; Fig. 2C), although this effect was reliably stronger in the Imposed selection than in the Free selection condition (*F*_1,29_ = 6.20, *p* = 0.019). Apparently, some selection conflict may have occurred even in the Free selection condition. Finally, if the observed conflict signal was instrumental in bringing about slower saccades towards changed targets, this should be reflected in a correlation between theta power and switch costs. Indeed, stronger theta power for switch compared to repeat trials predicted higher switch costs across observers in the Imposed selection condition (robust %-bend correlation *r* = 0.53, *p* = 0.002; Fig. 2D), but not in the Free selection condition (*r* = 0.17, *p* = 0.37). Moreover, the condition differences in switch-related frontal theta correlated positively with condition differences in saccadic switch-costs (*r* = 0.47, *p* = 0.009). Beta-suppression did not correlate with behavior in either of the two conditions (all *p* > 0.55).

Taken together, the above results uncover a clear qualitative dissociation between proactive, preparatory switching reflected in posterior parietal and sensorimotor beta-band suppression, and reactive, conflict-related switching reflected in prefrontal theta-band enhancements. However, these results were obtained by pre-selecting EEG channels, and after standard trial-averaging methodology. To check whether these selections exhaustively captured all relevant mechanisms, we tested if we could predict at the single trial level whether an observer would switch or stay, based on the multivariate power distributions across the scalp for all time and frequency combinations. Linear-discriminant classifiers were trained on single-trial topographic distributions across the entire scalp, in dissociating switch from repeat “classes” across time and frequency (32). The performance of these classifiers was then trained on the same time-frequency points (through a cross-validation procedure; see Methods). In a time-window starting 500 ms prior to the anticipated search displays in the Free selection condition, a cluster of activity comprising the alpha-to-beta band (10-30 Hz) indeed predicted whether in the *upcoming* trial, the saccade was going to be directed towards a different (switch) or same (repeat) target as in the previous trial (*p* < 0.001; Fig. 3A). This effect re-appeared after the saccade (~500 ms post-stimulus), in a slightly lower frequency range (6–23 Hz; *p* < 0.001). Classification accuracy in the Imposed selection condition showed a similar broadband post-stimulus increase (*p* < 0.001). However, and crucially, when target selection was task-imposed there was no pre-stimulus beta activity that was predictive of target switches; instead, the significant post-stimulus classification cluster contained relatively stronger modulations in the lower frequency range of delta-theta (2–8 Hz). We directly compared conditions in these two time windows and frequency bands, and found that, as expected, the Free selection condition showed stronger beta decoding prior to the search display (*t*_29_ = 5.61, p < 0.001; Fig. 3B), whereas the Imposed selection condition showed stronger theta decoding during search (*t*_29_ = 2.76, p < 0.010; frequency by condition interaction: *F*_1,29_ = 30.55, *p* < 0.001). The forward-transformed topographies of the classifier weights served as a further validation of our initial channel selection. Such “class-separability maps” are equivalent to the univariate difference between conditions (33) (see methods). These maps confirmed that pre-stimulus decoding in beta (10–30 Hz) was reflected in a suppression over midfrontal-premotor channels during Free selection (*p* = 0.002; Fig. 3C), while post-stimulus decoding in theta (2–8 Hz) was reflected in a frontoparietal enhancement under Imposed selection (*p* < 0.001).

**Fig. 3.**
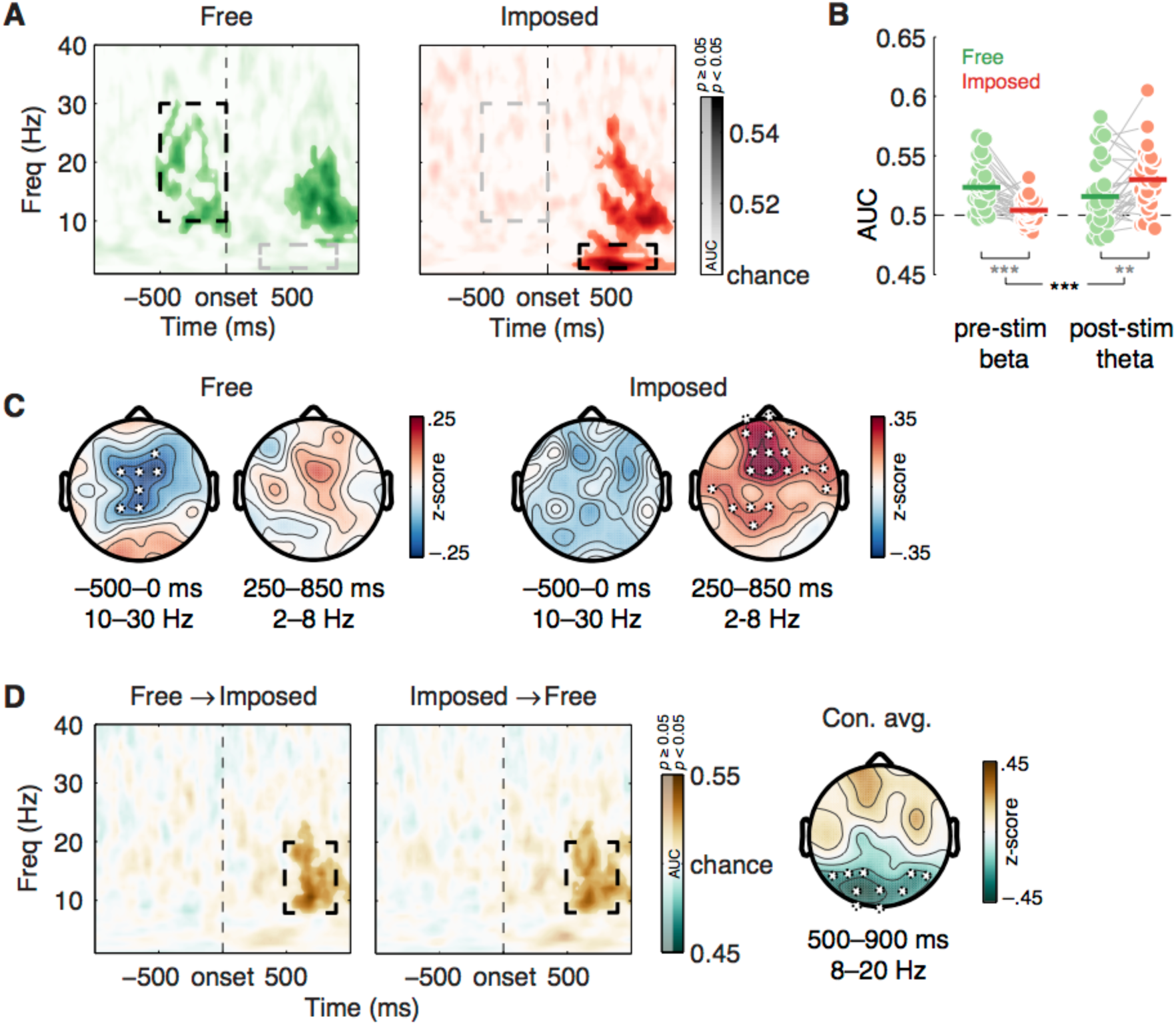
Multivariate classification findings. A) Classification accuracy (as measured through the area under the curve; AUC) after training a classifier on scalp-patterns of power at each time-frequency point in the Free (left; green) and Imposed (right; red) selection condition, using single-trial labels of ‘switch’ and ‘repeat’, and testing the classifier weights on the same time-frequency points through cross-validation. Saturated colors show significant clusters of AUC after cluster-size thresholding at p < 0.05. Dashed black/grey rectangles demarcate the time-frequency windows used to display weight topographies in B. B) Classifier accuracy per individual observer, showing direct comparisons between the Free and Imposed selection conditions. AUC values were averaged over parts of the significant time-frequency clusters shown in A, comprising the pre-stimulus alpha-to-beta band (10–30 Hz), and the post-stimulus delta-to-theta band (2–8 Hz). Each dot shows the accuracy of one individual observer. Horizontal lines show the group-average. Grey lines connecting the dots visualize the within-subject difference between Free selection and Imposed selection. ^∗∗^p < 0.01; ^∗∗∗^p < 0.001. C) Forward-transformed “class-separability”maps (33) of AUC within the time-frequency windows demarcated in A. Black-bordered white disks show clusters of significant neighboring electrodes after cluster-size thresholding at p < 0.01. D) Time-frequency maps of cross-condition decoding, after training on the Free selection condition and testing on the Imposed selection condition (left) and vice versa (center). Saturated colors highlight a common alpha-beta cluster late in the trial (~500–900 ms; p < 0.05) that generalized across the two conditions. Condition-average forward-transformed weights (right) of this common decoding effect showed a parieto-occipital scalp distribution (p < 0.01)

Post-saccade, the two conditions showed a comparable classification response in the alpha-beta range (Fig. 3A). Could this reflect a similar mechanism *after* a switch – whether freely chosen or task-imposed – has been initiated? Using cross-condition decoding (34), we tested whether the classifier weights trained on the data of one condition, could predict above chance whether trials from the other condition were switches or repeats. This analysis showed that the Free and Imposed selection conditions indeed cross-generalized to a common post-saccadic cluster in the alpha/lower-beta band, relatively late into the trial (8–20 Hz, 500–900 ms; *p* < 0.001; Fig. 3D). This signal may reflect the top-down implemented changes in prioritization, or ‘re-configured’ working memory content, in both conditions. Indeed, the condition-averaged forward-transformed weights in this time-frequency window showed a parieto-occipital suppression (p < 0.001), consistent with switches in priority states (35, 36).

## Discussion

Our study provides important new insights into the neural mechanisms underlying control of target selection in multiple-target search, . First, selecting an alternative target comes with specific neural state changes. Such state changes are not predicted by theories that claim that multiple-target search involves the equal, parallel prioritization of target representations, because a system that is prepared for both targets does not need to change its state. Instead, the data provide strong support for trial-by-trial priority shifts that can be traced through distinct neural signals.

Second, we provide the first evidence for dissociable control mechanisms over target selection. These mechanisms depend on whether the environmental context allows for free choice over which target to select or imposes such targets. We uniquely identified suppression of midfrontal/premotor beta-band (15–35 Hz) oscillatory activity as the signal that precedes free target switches with reduced behavioral cost. Importantly, time-frequency classifiers trained on single instances of beta-band scalp patterns could accurately predict these freely initiated switches about half a second before they actually happened. We therefore interpret it as the electrophysiological signature of proactive control. Consistent with this, when observers were forced to switch, no pre-stimulus beta suppression occurred. Instead, here a switch was followed by a transient burst of prefrontal delta/theta-band (2–8 Hz) oscillatory power, a signal that has been associated with reactive control, and which was positively correlated with individual differences in behavioral switch costs. We therefore interpret this signal as the electrophysiological signature of reactive control.

In task switching, proactive control has been linked to various event-related potential components in EEG, as well as fMRI BOLD modulations within a frontoparietal network (37-41). However, these studies used explicit cues instructing observers to switch stimulus-response mappings. Our findings provide a novel contribution also to this field, as we found anticipatory beta suppression to precede an endogenous switch under free choice. This is in line with several findings linking beta-band dynamics to represent the current sensorimotor or cognitive state in perceptual decision making and working memory (28, 42-46). Primate studies have also shown beta band involvement in the top-down maintenance of search targets (47, 48). Conversely, beta activity has been proposed to be inversely related to an upcoming voluntary change-of-action (49), including the amount of free choice a monkey is allowed in choosing the order in which it fixates multiple targets (50). Finally, the midfrontal topography of the signal is consistent with fMRI studies implicating the medial frontal cortex in self-initiated behavior (20, 23, 24).

Theta oscillations on the other hand have been proposed to constitute a key mechanism in medial and lateral prefrontal cortex that detects conflict between competing response alternatives (27), and implements cognitive control (31). As such they serve behavioral adjustments, which typically results in slower performance (51, 52). Here, we witnessed a qualitatively similar prefrontal theta-band modulation triggered by what may analogously be an internal conflict between competing target representations. Alternatively, task-imposed switching between representations may involve novelty processing, the manipulation of working memory content, or can be considered a form of prediction error, all of which have also been shown to elicit changes in prefrontal theta, with occasional extensions to the delta range (26, 27, 53-57). Interestingly, endogenous free switches also showed a similar but weaker theta increase in the trial-averaged power analysis, which did not correlate with individual switch-costs (and did not emerge in the multi-variate analysis). It is likely that the very presence of both template-matching targets in the search display invokes some competition, which needs to be resolved even during free proactive selection.

Note that although the current data support the claim that changing target selection is accompanied by changes in the observer’s priority state, they do not support the stronger claim that only a single target representation is active – and can thus be actively looked for – at a time (10). The data are also consistent with multiple representations being active in parallel, but with a measurable difference in attentional weights (8, 36) or priming (9), leading to various degrees of priority of different target representations. What our results indicate is that this differential weighting is disruptive when observers have no choice over which target they will encounter, but can be overcome when the observer is given full proactive control.

To conclude, we provide the first direct support for two modes of control being operative in multiple-target search, which are expressed in widespread qualitative differences in the time-frequency landscape of the EEG signal. Moreover, these signal patterns are predictive of when a switch will occur (under proactive control conditions), or what the switch cost will be (under reactive control conditions). Our study not only bridges different findings within the field of visual search, but also connects concordant ideas in the fields of attention and cognitive control.

## Materials and Methods

### Participants

Thirty healthy human participants (18 male) with normal or corrected-to-normal vision participated in this study for course credit or monetary compensation. The study was conducted in accordance with the Declaration of Helsinki and was approved by the faculty’s Scientific and Ethical Review Board (VCWE). Written informed consent was obtained.

### Task

Participants performed two conditions of a multiple-target gaze-contingent visual search task (15), in a blocked-design. The two versions differed in whether both or only one of two memorized search targets were available for selection in the subsequent search displays. The following task settings and stimulus parameters were identical across these two conditions.

The stimulus set consisted of six colored disks extending over a visual angle of 1.3°. The RGB-values of these colors were (0, 128, 175) for blue, (196, 79, 104) for red, (79, 123, 51) for green, (163, 107, 34) for brown, (142, 101, 183) for purple, and (120, 120, 120) for gray. All colors were isoluminant (M = 20 cd/m2). The background color was black (0, 0, 0).

After fixation drift correction (see section “Apparatus and eye tracking”), a block began with a fixation cross for 500 ms, followed by a cue display for 2500 ms and another fixation cross for 500 ms (see Figure 1). The cue display consisted of two colored disks 1.06° to the left and right of fixation and indicated the two target colors for the upcoming sequence of 40 search displays. The search displays each consisted of four colored and two identical gray disks, arranged in a hexagonal lattice with vertical rows and each at a distance of 3.9° from the hexagon’s center, which coincided with the fixation cross. Because of their regular positioning within a hexagon, the complete lattice on which stimuli could appear resembled a honeycomb structure. Participants were instructed to make a single eye movement towards a disk that matched either one of the target colors. The other items were distractors, not to be fixated. After target fixation, the stimuli were removed from the display and the fixated target was replaced by a white, filled circle, spanning 0.2°, to provide participants with a fixation point while participants waited for the next search display. If gaze position was not further than 1.95° away from this fixation point, the next display appeared after 850 to 1050 ms (randomly jittered). If participants failed to fixate this point for 5000 ms, a warning message appeared in the middle of the screen for 1000 ms, reminding them to look at the fixation point while waiting for the next search display. Because the coordinates of the previously fixated target determined the position starting point for the next display (the center of the hexagonal lattice), the search moved across the screen throughout a block, resembling natural eye movements during visual search when all items are present simultaneously. When the stimulus sequence approached an edge or the corner of the screen, the target (or targets) were randomly assigned to one (or two) of the three positions in the hexagon that were closest to the center of the screen, such that the next fixation would be directed away from the edge or corner. Although in such case the number of positions at which the targets could appear was thus reduced, participants still could not predict where exactly a specific color would appear.

Fixations had to land within a 2° visual angle radius around the target to be counted as valid. This ensured that fixations for targets and/or distractors could never overlap. If participants fixated one of the distractors, they received auditory feedback and were required to make a corrective eye movement towards a target. The search was aborted if no target was fixated within 3000 ms, and a new search display appeared.

There were two main factors. First, at the block level, target availability was manipulated. In the Free selection condition, both cued target colors appeared in the search display together with two gray and two colored distractors. In contrast, in the Imposed selection condition, only one of the two cued colors appeared in the search display together with two gray distractors and one colored distractor. Note that distractor colors remained fixed at the block level, and could be target colors in other blocks. The second factor was whether target color selection switched or repeated. Note that this latter factor was determined by either the observer (Free selection condition), or by a random sampling procedure, in which a sequence of target switches and target repetitions was randomly drawn (with replacement) from a pool of potential sequences (Imposed selection condition). Note that only the sequence of target switches and repeats was replayed, not the specific colors or positions of the search items, so that participants could not anticipate where a particular search target would appear. In order to match switch rate and streak length (successive switch or repeat trials) between conditions, sequences that were obtained during Free selection blocks were used to constitute the pool of replay sequences for Imposed selection blocks, for each participant separately. The pool of replay sequences to draw from would grow as the experiment progressed. Because at the outset of the experiment we did not have any sequences yet to fill the pool with, we initialized the pool with four pre-specified random sequences of target switches and repetitions (one each for six, eight, ten and twelve switches per block). Having a small proportion of fully random sequences also further prevented participants from recognizing the order of switches and repetitions in the sequences, while still closely matching switch rates between conditions. A paired sample t-test showed only a marginal (non-significant) difference between switch rates in the two conditions (*t*_29_ = 1.91, *p* = 0.07; Free selection: 31.2%, Imposed selection: 28.9%). As a double-check, we also asked participants after the experiment whether they were aware of this replay manipulation in the Imposed selection blocks, and none of them were.

In total, there were 40 blocks consisting of 40 search displays each. The five potential target colors were combined into 10 unique two-color cue combinations. Per target availability condition, each of these combinations was used twice as the pair of target colors for a block. Before the experiment started, observers practiced two blocks of both the Free and Imposed selection conditions.

### Apparatus and eye tracking

The experiments were designed and presented using OpenSesame (v3.1.4), (58) in combination with PyGaze (v0.6), an eye-tracking toolbox (59). Stimuli were presented on a 22 inch (diagonal) Samsung Syncmaster 2233RZ with a resolution of 1680 x 1050 pixels and refresh rate of 120 Hz at a viewing distance of 75 cm. Eye movements were recorded with the SR Research Eyelink 1000 tracking system (SR Research Ltd., Mississauga, Ontario, Canada) at a sampling rate of 1000 Hz and a spatial resolution of 0.01° visual angle. The experiments took place in a dimly lit, sound-attenuated room. The experimenter received real-time feedback on system accuracy on a second monitor located in an adjacent room. After every block, eye-tracker accuracy was assessed, and improved as needed by applying a 9-point calibration and validation procedure.

### EEG recording and cleaning

Concurrently with the eye-tracking (ET) data, electroencephalogram (EEG) data were acquired at 512 Hz from 64 channels (using a BioSemi ActiveTwo system; Amsterdam, the Netherlands) placed according to the international 10-20 system, and from both earlobes (used as reference). Offline, EEG and ET data were first co-registered using the EYE-EEG toolbox (v0.4; (60)) for EEGLAB (v12.0.2.3b; (61) in Matlab (2014a and 2015a; The Mathworks). All standard settings of the EYE-EEG tutorial were used (see www2.hu-berlin.de/eyetracking-eeg); the minimum plausible interval between saccades was set to 50 ms; from clusters of saccades within this interval, only the first was stored. Quality of EEG-ET synchronization was visually inspected using recommendations from the EYE-EEG tutorial; all data sets showed good synchronization and eye movement properties (i.e. fixation heatmaps and saccade angular histograms).

Next, EEG data were high-pass filtered at 0.5 Hz prior to time-frequency analysis to remove drifts and other non-stationarities (62), and 2.5 Hz solely for ICA to improve its signal-to-noise ratio (63, Dimigen, personal communication). Continuous EEG data were epoched from –2.5 to 3 s surrounding the onset of the search display (to avoid edge artifacts resulting from wavelet filtering, see below). The vertical electro-oculogram (VEOG) was recorded from electrodes located 2 cm above and below the right eye, and the horizontal EOG (HEOG) was recorded from electrodes 1 cm lateral to the external canthi. The EOG data were used together with EEG and ET data for automatic detection of oculomotor independent components (see below). Epochs were baseline-normalized using the whole epoch as baseline, which has been shown to improve independent component analysis (ICA; (64). Prior to cleaning, the data were visually inspected for malfunctioning electrodes, which were temporarily removed from the data (17 out of 30 participants had 1–3 malfunctioning electrodes).

To detect epochs that were contaminated by muscle artifacts, we used an adapted version of an automatic trial-rejection procedure as implemented in the Fieldtrip toolbox (65), using a 110 and 140 Hz pass-band to capture high-frequency muscle activity, and allowing for variable z-score cut-offs per participant based on the within-subject variance of z-scores. This procedure resulted in an average of 7.86% rejected trials (min–max across participants: 1.56–21.38%). After trial rejection, we performed ICA as implemented in EEGLAB only on the clean EEG, and EOG electrodes. Next, correlations between ET and independent components were used to automatically detect oculomotor artifacts, using the variance-ratio criterion suggested by (66) and as implemented in EYE-EEG; we removed on average 3.63 components (min–max across participants: 1–5). Finally, the malfunctioning electrodes identified before ICA were interpolated using EEGLAB’s `eeg_interp.m` function.

We only selected those trials that had a “clean” saccade-trajectory from fixation after search-display onset, to final fixation on a target-matching disk (which marked search-display offset). That is, intermediate fixations within such a trajectory had to fall within 30° around a straight line from initial fixation to (one of two) target(s). Trials that did not meet this criterion may have had trajectories in which saccades were first drawn towards distractors, even though they finally landed on a correct target. This selection procedure together with EEG artifact rejection resulted in an average of 360 (min–max across participants: 142–599) repeat and 145 (28–316) switch trials in the Free selection condition; Imposed selection condition: 351 (177–521) repeat and 113 (45–188) switch trials.

For time-frequency analyses, first the surface Laplacian of the EEG data was estimated (67, 68), which sharpens EEG topography and filters out distant effects that may be due to volume conducted activity from deeper brain sources (69, 70). The Laplacian can thus be interpreted as a spatial high-pass filter. For estimating the surface Laplacian, we used a 10th-order Legendre polynomial, and lambda was set at 10^−5^.

### Behavioral analysis

Our main behavioral variable of interest was trial-averaged latencies of the first eye movement (dwell time before a saccade toward a target was executed). Mean saccade latencies were computed separately for the Free and Imposed selection blocks, and separately for repeat trials (selected target color at trial N was the same as the selected target color at trial N-1) and switch trials (selected target color at trial N was different from the selected target color at trial N-1). We took the first saccade after search display onset, provided that it met the selection criterion as described above. Next, a saccade latency filter was applied, in which saccades quicker than 100 ms and slower than 3 SDs above the block mean for that participant were excluded (average of 2.5% of all trials). Average saccade latencies per participant were entered in a repeated measures ANOVA with factors trial type (repeat and switch) and condition (Free and Imposed), using JASP (Version 0.9; jasp-stats.org).

### EEG Time-frequency decomposition

Epoched EEG time series were decomposed into their time-frequency representations with custom-written MATLAB code (github.com/joramvd/tfdecomp). Each epoch was convolved with a set of complex Morlet wavelets with frequencies ranging from 1 to 40 Hz in 50 linearly spaced steps. Wavelets were created by multiplying perfect sine waves (*e*^*i*2*πft*^, where *i* is the complex operator, *f* is frequency, and *t* is time) with a Gaussian (*e*^−*t*^2^/2*s*^2^^, where *s* is the width of the Gaussian). The width of the Gaussian was set according to *s* = *δ*/2*πf*, where *δ* represents the number of cycles of each wavelet, linearly spaced between 3 (for 1 Hz) and 12 (for 40 Hz) in order to have a good trade-off between temporal and frequency precision. From the complex convolution result *Z_t_* (down-sampled to 40 Hz), an estimate of frequency-specific power at each time point was defined as [real(*Z_t_^2^*) + imag(*Z_t_^2^*)]. Single-trial power at each time-frequency point was used for a linear discriminant classification analysis (see below). Trial averaged power at each time-frequency point was decibel normalized according to 10^∗^log10(power/baseline), where for each channel and frequency, the condition averaged power during the entire trial served as baseline activity. We chose this baseline procedure because in a fast-paced saccade-driven trial design there is no optimal neutral baseline time window in e.g. the inter-trial interval, because of potential condition differences in both pre-stimulus and pre- and post-saccadic activity. Some baseline normalization procedure is nonetheless necessary to transform frequency-specific power to one common scale (i.e. to remove the 1/f scaling of power), and to correct for single-trial outliers (raw power cannot go below zero but can take relatively large values). Importantly, our main dependent variable was the difference in time-frequency power between switch and repeat trials (switch > repeat), in which any common deviation from “baseline” was subtracted out.

### EEG Multivariate pattern analysis

In addition to univariate time-frequency analysis on each single electrode, we applied a backward decoding classification algorithm (linear discriminant analysis or LDA) with all 64 channels as features and ‘switch’ and ‘repeat’ as classes, on time-frequency decomposed power. The goal of this analysis was to test whether a classifier could learn from spatial patterns of power modulations in specific time-frequency intervals, whether a participant at a single trial was going to repeat target selection or switch to a different target, and whether this would differ between the Free and the Imposed selection condition. Moreover, given that we used pre-stimulus activity, we could also test whether classifiers could predict such a choice before it happened.

For this analysis we used ADAM (the Amsterdam Decoding and Modeling toolbox http://www.fahrenfort.com/ADAM.htm), a freely available Matlab toolbox for backward decoding and forward encoding modeling of EEG and MEG data (32), replacing the standard time-frequency decomposition algorithm in that toolbox with our custom written time-frequency decomposition. Training and testing was done on the same data, for the Free and Imposed selection condition separately, using a 10-fold cross-validation procedure: first, trials for each of the two conditions were randomized in order, and divided into 10 equal-sized folds; next, a leave-one-out procedure was used on the 10 folds, such that the classifier was trained on 9 folds and tested on the remaining fold, and each fold was used once for testing. Classifier performance was then averaged over folds. Because there were more repeat than stay trials, we balanced the two classes through over-sampling, to ensure that during training the classifier would not develop a bias for the overrepresented class (see (32) for details). The classification performance output metric was the Area Under the Curve (AUC), with the curve being the receiver operating curve of the cumulative probabilities that the classifier assigns to instances as coming from the same class (true positives) against the cumulative probabilities that the classifier assigns to instances that come from the other class (false positives). AUC takes into account the degree of confidence (distance from the decision boundary) that the classifier has about class membership of individual instances, rather than averaging across binary decisions about class membership of individual instances (as happens when computing standard accuracy). As such the AUC is considered a sensitive, nonparametric and criterion-free measure of classification (71). We also inspected the spatial distribution of the classifier weights, through the product of the classifier weights and the original data covariance matrix at each time-frequency point. This procedure results in “class separability maps”, and are equivalent to the topographical maps of univariate difference between classes (33), although now numerically resulting from a decoding analysis.

### Statistics

Statistical analyses were done using group-level permutation testing with cluster correction (72). For decibel-normalized power, this was done on a switch > repeat contrast, for the Free and Imposed selection conditions separately, and on the double contrast of Free (switch > repeat) > Imposed (switch > repeat). For the multivariate pattern analysis results, this was done on AUC values above chance (0.5) for Free and Imposed selection separately. In all permutation tests, group-level *t*-values were first computed for the above contrasts, and for every time-frequency point. These *t*-values were thresholded at *p* < 0.05, yielding clusters of significant time-frequency power modulations. The *t*-values in each of these observed clusters were summed. Next, in 2000 iterations, the condition labels (e.g. power values of switch versus repeat, or the AUC value and its 0.5 reference value) were randomized for each subject, prior to performing *t*-tests on these permuted data. The sum of *t*-values within the largest cluster was saved into a distribution of summed cluster *t*-values. This distribution reflected cluster-level effect-sizes under the null-hypothesis of no effect. Finally, observed clusters with summed t-values smaller than the 95th percentile of the null-distribution (corresponding to *p* ≥ 0.05) were removed. This approach ensures a correction for multiple comparisons by taking into account clusters of spuriously significant time-frequency points that occur purely by chance. Time-frequency cluster tests were done on the average activity of electrodes FC1, FCz and FC2, which we selected based on previous findings from our lab (73). To visualize the spatial distribution of resulting time-frequency clusters, we next averaged over the activity within these clusters, and tested the same trial-type and condition contrasts over all channels, using cluster-correction across space instead of time-frequency, now with a (pre-)cluster-threshold of *p* < 0.01. To evaluate candidate clusters, we used Fieldtrip’s `neighbours` structure for 64-channel Biosemi layout, and we set 1 channel as the smallest possible cluster. All reported “cluster-corrected” *p*-values in the Results section refer to proportion of permuted clusters under the null hypothesis that were larger than the observed cluster.

Additionally, we ran parametric paired samples t-tests and repeated measures ANOVA’s with factors Condition (Free, Imposed) and Trial type (switch, repeat) over highlighted time-frequency windows, using JASP (version 0.9; jasp-stats.org). We further tested for cross-subject correlations between brain and behavior measures using the robust percentage-bend correlation metric that de-weights outliers (30).

## Acknowledgements

Author contributions: J.v.D., C.N.L.O., E.O. and J.J.F. designed research; J.v.D. and E.O. performed research; J.v.D analyzed data; J.v.D, J.J.F., E.O., and C.N.L.O. wrote the paper. J.J.F. and C.N.L.O share senior authorship. This work was supported by Consolidator Grant number ERC-2013-CoG-615423 of the European Research Council, awarded to C.N.L.O. The authors declare no conflict of interest. We thank Grace Pulsford for assistance in data collection. We thank Olaf Dimigen for his advice on preprocessing EEG combined with eye tracking.

## References

1. Liu T, Jigo M (2017) Limits in feature-based attention to multiple colors. Att Percept Psychophys 79(8):2327–2337.

2. Houtkamp R, Roelfsema PR (2008) Matching of visual input to only one item at any one time. Psychol Res 73(3):317–326.

3. Grubert A, Eimer M (2013) Qualitative differences in the guidance of attention during single-color and multiple-color visual search: Behavioral and electrophysiological evidence. J Exp Psychol Hum Percept Perform 39(5):1433–1442.

4. Mitroff SR, Biggs AT, Cain MS (2015) Multiple-target visual search errors: Overview and implications for airport security. Policy Insights Behav Brain Sci 2(1):121–128.

5. Menneer T, Cave KR, Donnelly N (2009) The cost of search for multiple targets: effects of practice and target similarity. J Exp Psychol Appl 15(2):125–139.

6. Maljkovic V, Nakayama K (1994) Priming of pop-out: I. Role of features. Mem Cognit 22(6):657–672.

7. Dombrowe I, Donk M, Olivers CNL (2011) The costs of switching attentional sets. Att Percept Psychophys 73(8):2481–2488.

8. Found A, Müller HJ (1996) Searching for unknown feature targets on more than one dimension: Investigating a “dimension-weighting” account. Percept Psychophys 58(1):88–101.

9. Kristjánsson Á, Campana G (2010) Where perception meets memory: a review of repetition priming in visual search tasks. Att Percept Psychophys 72(1):5–18.

10. Olivers CNL, Peters J, Houtkamp R, Roelfsema PR (2011) Different states in visual working memory: when it guides attention and when it does not. Trends Cogn Sci 15(7):327–334.

11. Beck VM, Hollingworth A, Luck SJ (2012) Simultaneous control of attention by multiple working memory representations. Psychol Sci 23(8):887–898.

12. Beck VM, Hollingworth A (2017) Competition in saccade target selection reveals attentional guidance by simultaneously active working memory representations. J Exp Psychol Hum Percept Perform 43(2):225–230.

13. Grubert A, Eimer M (2015) Rapid parallel attentional target selection in single-color and multiple-color visual search. J Exp Psychol Hum Percept Perform 41(1):86–101.

14. Kristjánsson T, Thornton IM, Kristjánsson Á (2018) Time limits during visual foraging reveal flexible working memory templates. J Exp Psychol Hum Percept Perform 44(6):827–835.

15. Ort E, Fahrenfort JJ, Olivers CNL (2017) Lack of free choice reveals the cost of having to search for more than one object. Psychol Sci 28(8):1137–1147.

16. Ort E, Fahrenfort JJ, Olivers CNL (2018) Lack of free choice reveals the cost of multiple-target search within and across feature dimensions. Att Percept Psychophys 10:104–14.

17. Monsell S (2003) Task switching. Trends Cogn Sci 7(3):134–140.

18. Braver TS (2012) The variable nature of cognitive control: a dual mechanisms framework. Trends Cogn Sci 16(2):106–113.

19. Geng JJ (2014) Attentional mechanisms of distractor suppression. Curr Dir Psychol Sci 23(2):147–153.

20. Taylor PCJ, Rushworth MFS, Nobre AC (2008) Choosing where to attend and the medial frontal cortex: an FMRI study. J Neurophysiol 100(3):1397–1406.

21. Forstmann BU, Ridderinkhof KR, Kaiser J, Bledowski C (2007) At your own peril: an ERP study of voluntary task set selection processes in the medial frontal cortex. Cogn Affect Behav Neurosci 7(4):286–296.

22. Frith CD, Haggard P (2018) Volition and the brain - revisiting a classic experimental study. Trends Neurosci 41(7):405–407.

23. Wisniewski D, Reverberi C, Tusche A, Haynes J-D (2015) The neural representation of voluntary task-set selection in dynamic environments. Cereb Cortex 25(12):4715–4726.

24. Schuck NW, et al. (2015) Medial prefrontal cortex predicts internally driven strategy shifts. Neuron 86(1):331–340.

25. Cunillera T, et al. (2011) Brain oscillatory activity associated with task switching and feedback processing. Cogn Affect Behav Neurosci 12(1):16–33.

26. Cavanagh JF, Zambrano Vazquez L, Allen JJB (2011) Theta lingua franca: A common mid-frontal substrate for action monitoring processes. Psychophysiology 49(2):220–238.

27. Cohen MX (2014) A neural microcircuit for cognitive conflict detection and signaling. Trends Neurosci 37(9):480–490.

28. Donner TH, Siegel M (2011) A framework for local cortical oscillation patterns. Trends Cogn Sci 15(5):191–199.

29. Helfrich RF, Knight RT (2016) Oscillatory dynamics of prefrontal cognitive control. Trends Cogn Sci 20(12):916–930.

30. Pernet CR, Wilcox R, Rousselet GA (2012) Robust correlation analyses: false positive and power validation using a new open source matlab toolbox. Front Psychol 3:606.

31. Cavanagh JF, Frank MJ (2014) Frontal theta as a mechanism for cognitive control. Trends Cogn Sci 18(8):414–421.

32. Fahrenfort JJ, van Driel J, van Gaal S, Olivers CNL (2018) From ERPs to MVPA Using the Amsterdam Decoding and Modeling Toolbox (ADAM). Front Neurosci 12:368.

33. Haufe S, et al. (2014) On the interpretation of weight vectors of linear models in multivariate neuroimaging. NeuroImage 87(C):96–110.

34. King JR, Dehaene S (2014) Characterizing the dynamics of mental representations: the temporal generalization method. Trends Cogn Sci 18(4):203–210.

35. de Vries IEJ, van Driel J, Olivers CNL (2017) Posterior α EEG dynamics dissociate current from future goals in working memory-guided visual search. J Neurosci 37(6):1591–1603.

36. de Vries IEJ, van Driel J, Karacaoglu M, Olivers CNL (2018) Priority switches in visual working memory are supported by frontal delta and posterior alpha interactions. Cereb Cortex 26:1513–15.

37. Sohn MH, Ursu S, Anderson JR, Stenger VA, Carter CS (2000) The role of prefrontal cortex and posterior parietal cortex in task switching. Proc Natl Acad Sci U S A 97(24):13448–13453.

38. Braver TS, Reynolds JR, Donaldson DI (2003) Neural mechanisms of transient and sustained cognitive control during task switching. Neuron 39(4):713–726.

39. Karayanidis F, et al. (2010) Advance preparation in task-switching: converging evidence from behavioral, brain activation, and model-based approaches. Front Psychol 1:25.

40. Rushworth MFS, Passingham RE, Nobre AC (2002) Components of switching intentional set. J Cogn Neurosci 14(8):1139–1150.

41. Astle DE, Nobre AC, Scerif G (2009) Applying an attentional set to perceived and remembered features. PLOS ONE 4(10):e7613.

42. Haegens S, et al. (2011) Beta oscillations in the monkey sensorimotor network reflect somatosensory decision making. Proc Natl Acad Sci U S A 108(26):10708–10713.

43. Donner TH, Siegel M, Fries P, Engel AK (2009) Buildup of choice-predictive activity in human motor cortex during perceptual decision making. Curr Biol 19(18):1581–1585.

44. Kloosterman NA, et al. (2015) Top-down modulation in human visual cortex predicts the stability of a perceptual illusion. J Neurophysiol 113(4):1063–1076.

45. Engel AK, Fries P (2010) Beta-band oscillations – signalling the status quo? Curr Opin Neurobiol 20(2):156–165.

46. Spitzer B, Haegens S (2017) Beyond the status quo: A role for beta oscillations in endogenous content (re)activation. eNeuro 4(4). doi:10.1523/ENEURO.0170-17.2017.

47. Buschman TJ, Miller EK (2007) Top-down versus bottom-up control of attention in the prefrontal and posterior parietal cortices. Science 315(5820):1860–1862.

48. Buschman TJ, Miller EK (2009) Serial, covert shifts of attention during visual search are reflected by the frontal eye fields and correlated with population oscillations. Neuron 63(3):386–396.

49. Jenkinson N, Brown P (2011) New insights into the relationship between dopamine, beta oscillations and motor function. Trends Neurosci 34(12):611–618.

50. Pesaran B, Nelson MJ, Andersen RA (2008) Free choice activates a decision circuit between frontal and parietal cortex. Nature 453(7193):406–409.

51. Kerns JG, et al. (2004) Anterior cingulate conflict monitoring and adjustments in control. Science 303(5660):1023–1026.

52. van Driel J, Swart JC, Egner T, Ridderinkhof KR, Cohen MX (2015) (No) time for control: Frontal theta dynamics reveal the cost of temporally guided conflict anticipation. Cogn Affect Behav Neurosci 15(4):787–807.

53. Barcelό F, Escera C, Corral MJ, Periáñez JA (2006) Task switching and novelty processing activate a common neural network for cognitive control. J Cogn Neurosci 18(10):1734–1748.

54. Munneke G-J, Nap TS, Schippers EE, Cohen MX (2015) A statistical comparison of EEG time- and time–frequency domain representations of error processing. Brain Res 1618:222–230.

55. Ullsperger M, Fischer AG, Nigbur R, Endrass T (2014) Neural mechanisms and temporal dynamics of performance monitoring. Trends Cogn Sci 18(5):259–267.

56. Nee DE, Kastner S, Brown JW (2011) Functional heterogeneity of conflict, error, task-switching, and unexpectedness effects within medial prefrontal cortex. NeuroImage 54(1):528–540.

57. Itthipuripat S, Wessel JR, Aron AR (2012) Frontal theta is a signature of successful working memory manipulation. Exp Brain Res 224(2):255–262.

58. Mathôt S, Schreij D, Theeuwes J (2011) OpenSesame: An open-source, graphical experiment builder for the social sciences. Behav Res 44(2):314–324.

59. Dalmaijer ES, Mathôt S, Van der Stigchel S (2013) PyGaze: An open-source, cross-platform toolbox for minimal-effort programming of eyetracking experiments. Behav Res 46(4):913–921.

60. Dimigen O, Sommer W, Hohlfeld A, Jacobs AM, Kliegl R (2011) Coregistration of eye movements and EEG in natural reading: analyses and review. J Exp Psychol Gen 140(4):552–572.

61. Delorme A, Makeig S (2004) EEGLAB: an open source toolbox for analysis of single-trial EEG dynamics including independent component analysis. J Neurosci Methods 134(1):9–21.

62. Cohen MX (2014) Analyzing Neural Time Series Data (MIT Press).

63. Winkler I, Debener S, Müller K-R, Tangermann M (2015) On the influence of high-pass filtering on ICA-based artifact reduction in EEG-ERP. Conf Proc IEEE Eng Med Biol Soc 2015:4101–4105.

64. Groppe DM, Makeig S, Kutas M (2009) Identifying reliable independent components via split-half comparisons. NeuroImage 45(4):1199–1211.

65. Oostenveld R, Fries P, Maris E, Schoffelen J-M (2011) FieldTrip: Open source software for advanced analysis of MEG, EEG, and invasive electrophysiological data. Comput Intell Neurosci 2011(1):156869–9.

66. Plöchl M, Ossandón JP, König P (2012) Combining EEG and eye tracking: identification, characterization, and correction of eye movement artifacts in electroencephalographic data. Front Hum Neurosci 6:278.

67. Perrin F, Pernier J, Bertrand O, Echallier JF (1989) Spherical splines for scalp potential and current density mapping. Electroencephalogr Clin Neurophysiol 72:184–187.

68. Kayser J, Tenke CE (2015) Issues and considerations for using the scalp surface Laplacian in EEG/ERP research: A tutorial review. Int J Psychophysiol 97(3):189–209.

69. Oostendorp TF, van Oosterom A (1996) The surface Laplacian of the potential: theory and application. IEEE Trans Biomed Eng 43(4):394–405.

70. Winter WR, Nunez PL, Ding J, Srinivasan R (2007) Comparison of the effect of volume conduction on EEG coherence with the effect of field spread on MEG coherence. Stat Med 26(21):3946–3957.

71. Hand DJ, Till RJ (2001) A simple generalisation of the area under the ROC curve for multiple class classification problems. Mach Learn 45(2):171–186.

72. Maris E, Oostenveld R (2007) Nonparametric statistical testing of EEG- and MEG-data. J Neurosci Methods 164(1):177–190.

73. van Driel J, Gunseli E, Meeter M, Olivers CNL (2017) Local and interregional alpha EEG dynamics dissociate between memory for search and memory for recognition. NeuroImage 149:114–128.

